# Fitness costs of parasitism depend on fine-scale density and resource availability in a wild ungulate

**DOI:** 10.64898/2026.04.07.716954

**Authors:** Adam Hasik, Alison Morris, Sean Morris, Katie Maris, Shane Butt, Amy R Sweeny, Josephine M Pemberton, Gregory F Albery

## Abstract

Resource competition and parasite exposure both present common density-dependent fitness costs for wild animals. Because launching effective immune responses is costly in terms of resources, parasites’ fitness costs should be further exacerbated in high-density, resource-depleted areas. To disentangle these relationships, we related density, parasitism, and resource availability to survival and fecundity across lifespan in a long-term study of wild red deer. All fitness measures declined with a combination of parasite count, greater density, and reduced resource availability. Beyond these relationships, as expected, local density and resource scarcity exacerbated survival costs of parasitism in calves, effectively undermining tolerance of infection. However, these synergistic relationships faded in yearlings and then reversed in adults, likely through age-structured selection biases. These findings emphasize that the costs of parasites and resource scarcity can be synergistic and intertwined with density in wild populations, accentuating the value of incorporating resource competition when examining parasite-dependent population regulation.

## Introduction

Population density is a fundamental driver of variation in fitness in wild animal populations. As density increases, individuals experience intensified competition for limited resources^1,2^ and increased exposure to parasites^3,4^, both of which can reduce host survival and reproduction. Parasites can play a central role in density-dependent population regulation, because higher host densities increase contact rates and environmental concentration of infectious stages, which then produces a growing fitness cost for each individual in the population^3-5^. A range of studies have documented positive relationships between aspects of host density and parasitism (or contact rates) in natural populations, supporting this possibility^6-9^. However, there are also a variety of more complex documented density-infection (or density-contact) trends, some of which could be explained by the fact that density determines not only parasites’ intensity and prevalence, but their fitness effects in space and time^10-12^. Nevertheless, density and parasites’ joint effects on host fitness are unclear in wild populations; specifically, their fine-scale distributions and interactions with resource availability have rarely been considered.

Resource availability affects a host’s nutritional state and immune defenses^13-15^, which influences both the ability to resist infection^16,17^ and tolerate its negative effects^18,19^. When resources are scarce, hosts may face myriad trade-offs between immunity and other physiological processes (e.g., reproduction), potentially exacerbating parasite infection or its fitness costs^20-22^. Conversely, abundant resources may allow hosts to compensate for infection or maintain effective immune responses, as is commonly demonstrated in the lab^23-26^. There is also some evidence for the same under field manipulations; for example, in wood mice (*Apodemus sylvaticus*) supplemented nutrition increases resistance to helminth infection^26^, but simultaneously increases host density, with the strongest effects occurring in the years of lowest baseline resources^27^. These relationships suggest that parasites may impose the greatest fitness costs where both infection risk and resource limitation are high. Critically, although both high parasite exposure and resource scarcity are possible outcomes of high density, they have never been linked with one another in the context of density dependence.

In cases where density drives greater exposure as well as greater vulnerability to infection, the impact of density-dependent parasite regulation could be even greater than often thought. That is, hosts at higher densities might not just be more likely to become infected, but to die once infected. Nevertheless, few studies have examined how these factors interact to influence fitness in natural populations. A particular challenge is that density, resource availability, and parasitism all vary spatially within and between populations^28^, but most studies of density dependence rely on population-level measures of density^29-31^, which may obscure substantial variation in the densities experienced by individual hosts^32^. Meanwhile, lab populations and experimental manipulations often struggle to replicate natural densities, levels of infection, and resource competition, particularly in concert with other important natural stressors. In wild and lab populations, ecological tolerance (which is related to, but distinct from, immunological tolerance) is quantified as the slope of fitness on parasitism^33-35^; variation in this slope represents variation in tolerance. To date, few studies have identified modifiers of tolerance in the wild, due largely to difficulties quantifying these slopes and their interactions with other factors.

Here, we investigate how density, resource availability, and parasitism jointly influence fitness in a long-term study population of red deer (*Cervus elaphus*) on the Isle of Rum, Scotland. This population has been monitored for five decades, providing detailed data on individual life histories, survival, reproduction, and behavior^36,37^. Since 2016, individuals have also been sampled to examine infection with environmentally-transmitted gastrointestinal helminths. Helminth infection intensity – specifically of strongyle nematodes (order: Strongylida) – varies seasonally and among individuals^38^, and previous work in this system has shown that parasite burdens can influence fitness of both juveniles^39,40^ and adults^41^ as well as being influenced substantially by reproductive investment and resource reallocation away from immunity^42^. Additionally, despite being less than 12km^2^ in area, there are strong spatial patterns of infection and immunity^43^, some of which are explained by variation in age and sociality^44^ as well as population density^8^. There is a strong spatial gradient of grazing availability, with abundant resources in the northern area balanced by greater density, and with a gradient of mortality towards the south arising from occasional shooting of individuals that stray outside the study area^36^. Overall, the strong spatial patterns of density and resource availability, combined with the demonstrated effect of resource allocation on parasite count, strongly imply that the population will exhibit density- and resource-dependent fitness costs of parasitism. We expect that population density will decrease fitness by reducing resource availability, and that these will synergistically exacerbate the costs of parasitism for fitness at multiple life stages.

## Methods

### Study system and quantification of infection

We collected data for this study from a population of individually-recognized red deer in the north block of the Isle of Rum, Scotland (57°N,6°20’W). Rum has a wet, mild climate and contains a mixture of high-quality (i.e., high nutrition) grassland and low-quality (i.e., high in tannins, harder to consume) blanket bog and heath. The study area runs ∼4km north to south and ∼3km east to west, totaling ∼12.7km^2^. In 1974 culling ceased in the study area, the population of females increased while that of males decreased; the population has been unmanaged and at carrying capacity since about 1980 (see Pemberton et al.^37^ for an in-depth history of the study population). The population is censused approximately every week for eight months of the year^36^ and currently contains ∼250 individuals, mainly consisting of females and their recent offspring.

We have collected data on the helminth parasite burden of the population since 2016 by non-invasively gathering fecal samples three times during the “deer year” (which runs May 1^st^-April 30^th^) in August (summer), November (autumn), and April (spring). The spring sampling occurs at the *end* of the deer year, when deer are in relatively poor condition at the end of winter. Observers note individually-recognized deer defecating from a distance and collect fecal samples without disturbing the deer. Samples are placed in plastic bags to keep them as anaerobic as possible and refrigerated at 4°C to prevent hatching or development of parasite propagules, with parasitological examination being conducted within three weeks to generate fecal egg counts per gram of feces (FECs) see^45^.

Many red deer are infected with strongyle nematodes (hereafter “strongyles”, a mix of different species with indistinguishable eggs). Strongyles are directly-transmitted parasites; infective stages contaminate vegetation via fecal pellets and are subsequently consumed by a new host^46^. Strongyle infections develop quickly such that calves excrete eggs within 2-3 months of birth, and the population exhibits strong seasonal variation in infection^38^. The highest counts are in April when the deer have just survived the winter and are in poorest condition, females are in the late stages of pregnancy, and green-up is only just starting. The lowest counts are in November when the deer (except males) are in good condition following their summer food intake and because the parasites themselves reduce their egg production during colder months.

For our analyses we calculated annual strongyle parasite burden by first log+1 transforming all values, after which we standardized strongyle counts within season such that the seasonal mean was 0 and the SD was 1 to account for the known seasonal variation.

### Measuring global and local density

We estimated the density experienced by each individual for each year of study using two metrics: population-level annual density (hereafter “global density”) and individual-level annual density (hereafter “local density”). Using local density more precisely represents within-population variation in the densities experienced by individual deer, which are associated with variation in parasite counts in this population^8^. Using global density allows us to understand if the previously documented population-level density-dependent relationships with deer survival and reproduction^1,47^ are still present after the population has been at carrying capacity for multiple decades.

We used census data for the years 2016-2024, when individuals’ identities and locations (to the nearest 100m) are recorded, to calculate both density metrics. Global density is simply a count of the number of adult females using the study area regularly during a given year. We calculated local density using a previously described pipeline for the study population^8,44^ and across other systems including sheep^10^ and reptiles^48^. This approach uses a kernel density estimator applied to all observations of each individual in each deer year. We took individuals’ annual centroids and fit a two-dimensional smooth to the distribution of the data, producing a two-dimensional spatial distribution of the population. Individuals were then assigned a local density value based on their location on this distribution.

### Measuring individual resource availability

To investigate whether resource availability influences fitness, we used a processed dataset of Landsat satellite-derived measures of the Normalized Difference Vegetation Index (NDVI). The dataset provides estimated annual maximum NDVI (NDVI_max_) across the study site at a pixel resolution of 30 x 30 m. We aggregated NDVI data to 100 x 100m squares, then overlaid deer sighting data onto the NDVI map and calculated mean NDVI_max_ values (hereafter NDVI) of pixels which overlapped with a sighting of each deer throughout a given year, giving us an estimate of the maximum amount of vegetation a given deer had access to during that year as in^8^. Full details can be found in the Supplemental Materials. NDVI values range from -1 to 1, with increasing positive values indicative of greater green biomass^49^. NDVI is widely-recognized as an accurate and reliable indicator of vegetation biomass and net primary productivity^50^, making it a suitable proxy of resource availability for herbivorous species. Specifically, links have been established between NDVI and either body mass or condition in arthropods^51^, mule deer^52^, roe deer^50^, and this population^8^, making it an important correlate of fitness for the deer in this study. Importantly, density and NDVI_max_ are negatively correlated in space, likely because the deer depress and remove resources (Fig. 1).

**Fig. 1.**
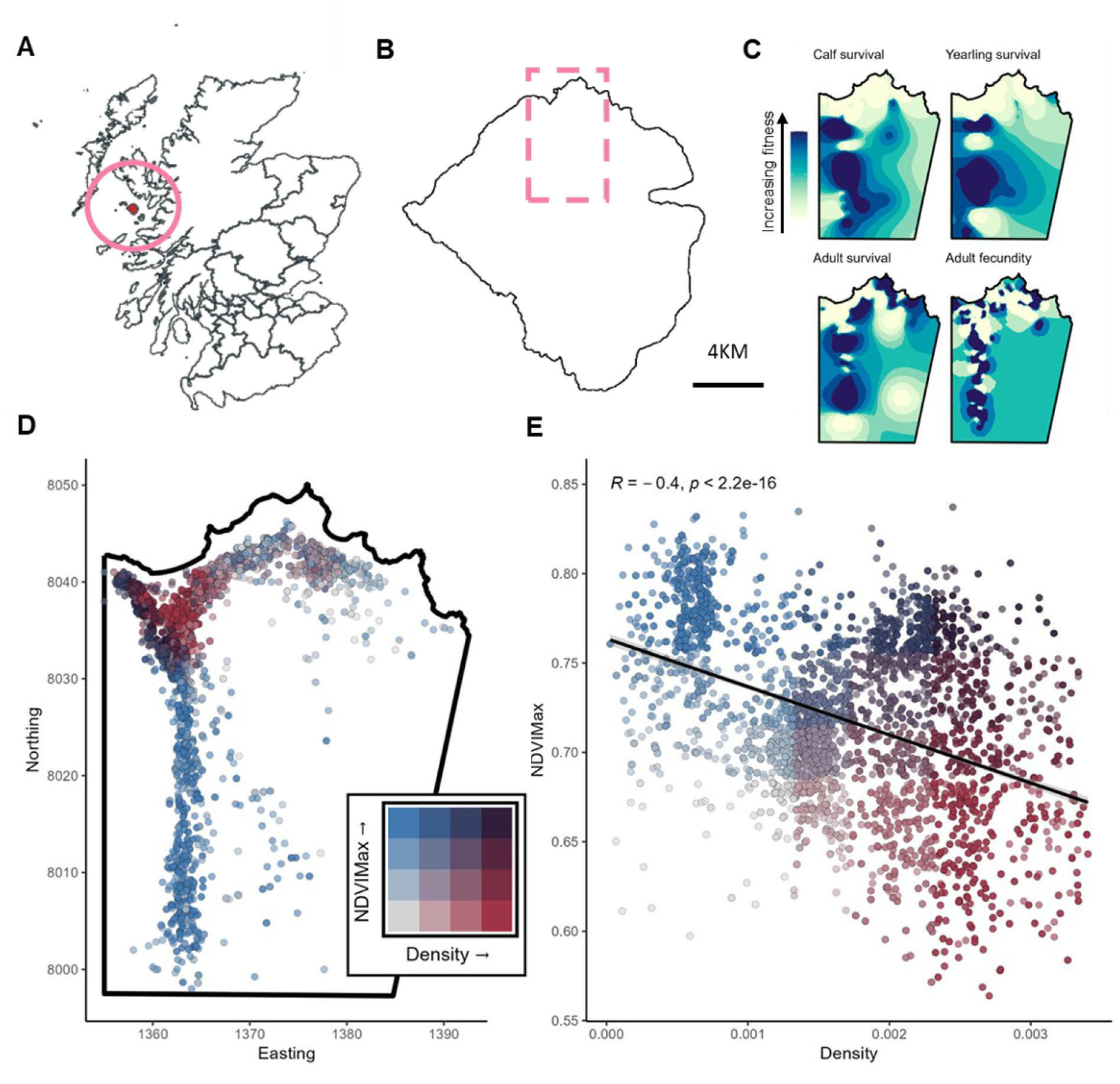
The distributions of local density, NDVI_max_, and fitness in the Rum deer population. A) The location of Rum within Scotland; B) the location of the study area within Rum. C) Spatial distributions of calf survival, yearling survival, adult survival, and adult fecundity. Taken from predictions from intercept-only spatial models with no other predictors, these are designed to depict the quantitative patterns of fitness across the population. Colors indicate predicted values on the response scale, with darker shades corresponding to higher predicted fitness. Across all fitness components, strong spatial structure is evident. D) Bivariate distribution of local population density and vegetation productivity (NDVI_max_) across the study population. Eastings and Northings are in units of 100m, so each tick is 10*100=1km. Points show individuals’ centroids colored according to their joint quantile classification of density (increasing to the right) and NDVI_max_ (increasing upwards), illustrating that high-density areas tend to coincide with areas of lower resource availability. This is confirmed by correlating the two in E), which depicts the same data, accompanied by a Pearson’s *R* calculation.

### Quantifying fitness components

We investigated overwinter survival of three age categories: calves (deer in their first year; *n*_*s*_ = 702 samples from *n*_*i*_ = 343 individuals), yearlings (in their second year; *n*_*s*_ = 405, *n*_*i*_ = 195), and adult females (aged 3 and above; *n*_*s*_ = 1509, *n*_*i*_ = 189). Survival is a binary variable, with 1 representing living through the winter into the start of the next deer year, and 0 representing death before the start of the next deer year. We removed individuals that were shot when they roamed outside of the study area from the analysis. In adult females, we also investigated fecundity in the following year (*n*_*s*_ = 1386; *n*_*i*_ = 169). Fecundity is also binary, with 1 for females that produced a calf in the next deer year, and 0 for those females that did not produce a calf in the next deer year. We used data for all individuals sampled in deer years 2016-2024, producing 1,386 individual-by-year combinations from 495 individuals across our four fitness model sets. All variables vary spatially (Fig. 1).

### Models relating density, NDVI, and parasitism to fitness

To understand how density, NDVI, and parasitism are associated with fitness variation, we constructed models for each of the response variables. We used Integrated Nested Laplace Approximation (INLA) models for all analyses to control for spatial autocorrelation in our data. INLA models are a deterministic Bayesian approach that allow for quantification of spatial effects and have been increasingly used for spatial analyses^28,43,53^. We fit all models in R v4.4.3^54^ using the R-INLA package^55,56^.

For each model set we first fit a base model including a variety of variables. These fell into three groups: (i) shared variables (global density, local density, NDVI, and parasite eggs per gram [EPG]); (ii) juvenile variables (sex, birth date, birth weight, and mother’s age); and (iii) adult variables (age and reproductive status). We calculated density metrics as described above. Reproduction status is a three-level factor based on established costliness^41,42,57^: “none” (did not reproduce); “summer” (calf died before October that deer year, generally thought to be relatively low cost) and “winter” (calf lived to October but died overwinter or lived through its first year, generally thought to be very costly). For the adult female survival model, we removed all females with the “summer” reproductive status before analysis, as they had a near-100% survival rate that prevented the model from fitting well. Sex is a two-level factor (female and male), while we scaled all continuous variables to have a mean of 0 and a standard deviation of 1.

We fit all models as binomial Generalized Linear Mixed Models (GLMMs), with random effects of ID and Year to account for pseudoreplication. We also included a random effect of maternal ID in the calf and yearling models. Finally, we added spatial random effects (i.e., SPDE) to all models using individuals’ annual centroids (mean Easting and Northing). The models were generally strongly improved by incorporation of the SPDE random effect (ΔDIC between 12.00 and 125.67); however, it did not qualitatively change any relationships, and we were interested in the influence of spatially-defined variables with which the SPDE would compete. As such, we focus on reporting the non-spatial models in all cases.

For each model set we fit the base model, which included either the juvenile or adult variables, and all of the shared variables. We then fit a series of interactions depending on the model, where all variables were allowed to interact with both log(EPG+1) as well as sampling season to investigate whether parasite-mediated relationships with fitness were contingent on the season in which the individual was sampled. We fit each interaction one at a time, and if any interaction reduced the deviance information criterion (DIC) by more than 2, the best-fitting was kept and the model addition was repeated. We approached the problem this way to avoid overloading the model, and because the predictors were quite well correlated and we were concerned about our ability to extricate these sources of variance. Ultimately, there was only one round of addition per model.

## Results

We discovered a range of relationships between density, resources, parasites, and fitness, both in juveniles (Fig. 2) and adults (Fig. 3). See Fig. 4 for the full model estimates.

**Fig. 2:**
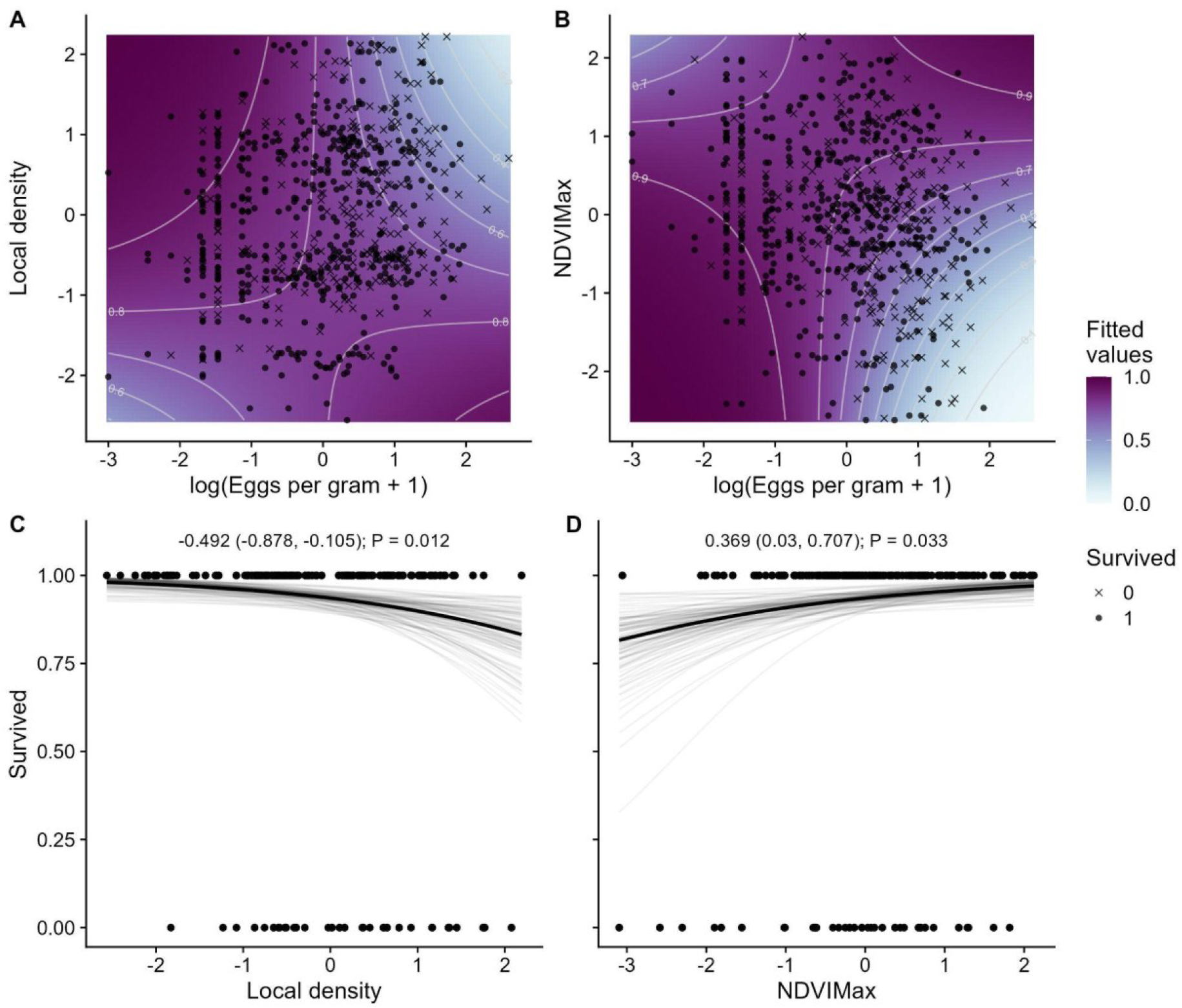
Model predictions of juvenile survival. (A–B) Predicted calf overwinter survival probability across gradients of parasite count (log[EPG + 1]) and (A) local density or (B) NDVI. Darker shading represents increased predicted probability of survival, delineated by grey contours and with proportional survival noted. Points show observed individual survival outcomes. (C-D) Relationships between (C) local density and (D) NDVI and yearling overwinter survival. Black lines show posterior mean predictions, grey lines show posterior samples, points indicate observed survival outcomes. Text above panels gives posterior mean estimates with 95% credible intervals and associated *p*-values. All continuous variables are scaled to have a mean of zero and standard deviation of 1.

**Fig. 3:**
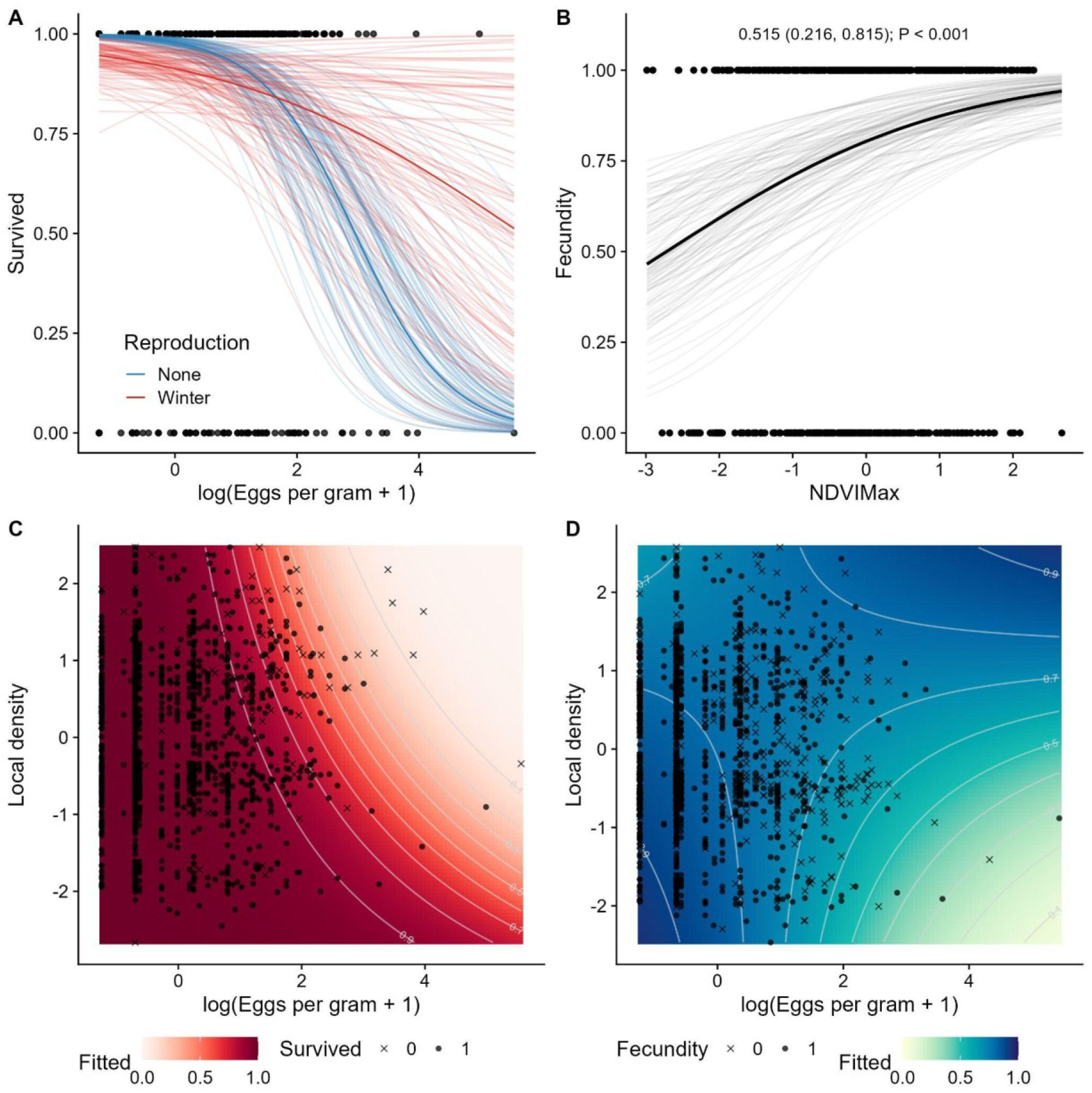
Model predictions of adult female fitness components. (A) Adult female overwinter survival probability decreases with increasing parasite count (log[EPG + 1]) differently for females whose calf survived to winter (red) versus non-reproductive females (blue). Transparent lines show posterior samples, opaque lines show posterior means, points show observed survival outcomes. (B) NDVI positively predicts adult female fecundity. The black line indicates posterior mean prediction, grey lines represent posterior samples, points indicate observed reproductive outcomes. Text provides posterior mean estimate with 95% credible interval and *p*-value. (C–D) Predicted adult female survival (C) and fecundity (D) across gradients of parasite burden and local density. Darker shading indicates higher predicted fitness values, delineated by grey contours and with proportional survival or fecundity noted; points indicate observed survival or reproduction outcomes. All continuous variables are scaled to have a mean of zero and standard deviation of 1.

**Fig. 4:**
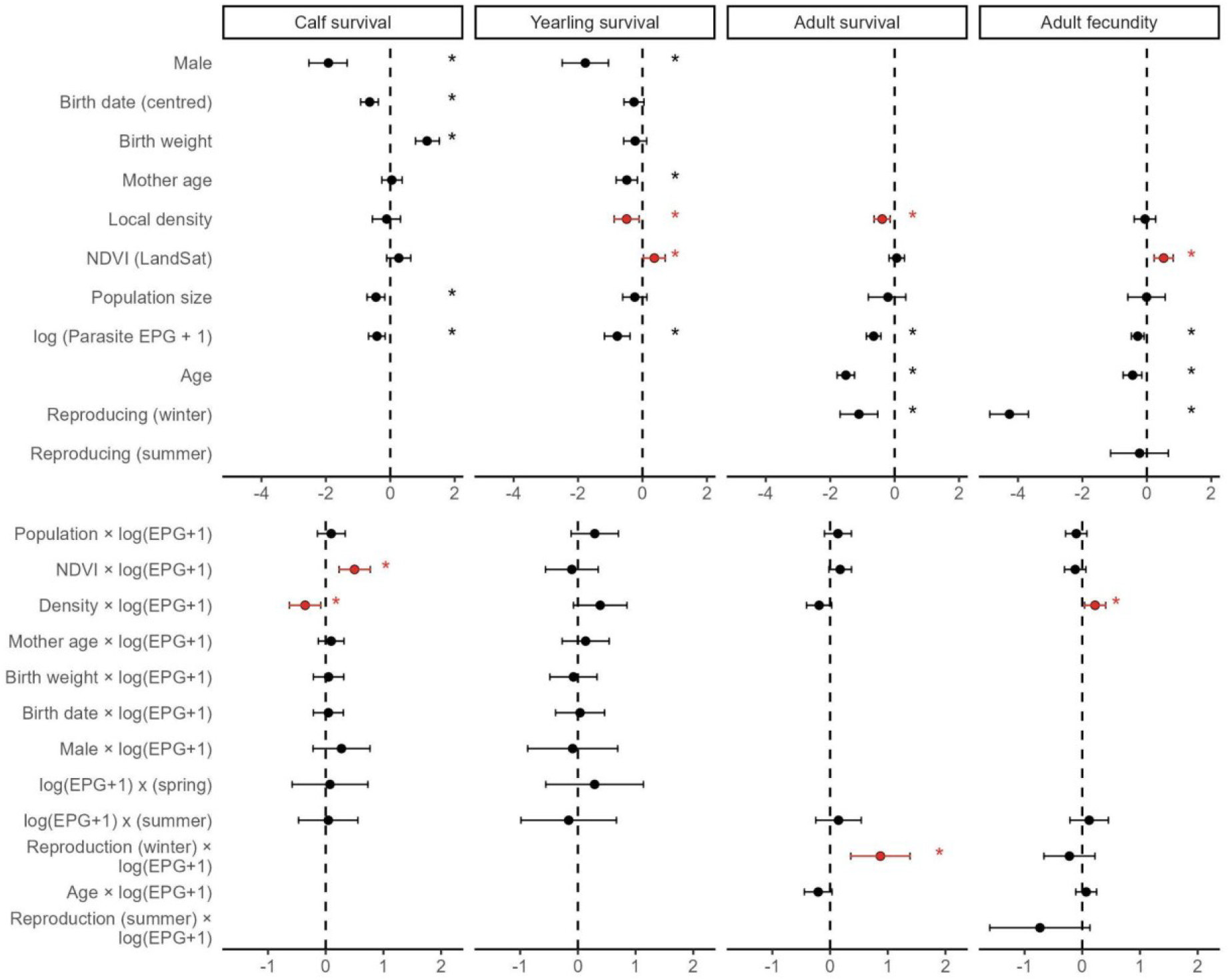
Forest plot depicting phenotypic relationships with fitness components. Posterior mean effect sizes (points) and 95% credible intervals (error bars) are taken from INLA models predicting (left to right) calf overwinter survival, yearling overwinter survival, adult female overwinter survival, and adult female fecundity. The top row shows models including base variables. The bottom row shows interaction terms between parasite burden and ecological or phenotypic predictors added on top of this model, one at a time. Asterisks denote effect sizes whose 95% credible intervals do not overlap zero. Red highlights have been used to identify findings in this system that are novel to this paper, all of which are depicted in Fig. 2-3.

As previously-demonstrated (reported with earlier references), calves were substantially less likely to survive following a high-density year (β = -0.44 (95% CI -0.70, -0.20), *p* = 0.001) and when exhibiting high parasite counts (β = -0.47 (95% CI -0.78, -0.22), *p* < 0.001). There were no direct relationships with NDVI or local density in calves: NDVI: (β = 0.22 (95% CI -0.11, 0.65), *p* = 0.29); local density: (β = -0.11 (95% CI -0.52, 0.30), *p* = 0.76); however, our analysis uncovered novel strong interactions between parasite counts and NDVI and density. Individuals with high parasite counts were much more likely to survive if they spent time in low-density (β = -0.34 (95% CI -0.59, -0.12), *p* = 0.01, Fig. 2a) or high-NDVI areas (β = 0.51 (95% CI 0.24, 0.76), *p* < 0.001, Fig. 2b). Both terms were significant when included individually, but only NDVI’s interaction remained significant when both were fit, presumably due to their relatively strong correlation. Finally, male, small, and late-born calves were also much less likely to survive (Fig. 4): male: (β = -1.97 (95% CI -2.58, -1.44), *p* < 0.001); birth weight: (β = 1.23 (95% CI 0.89, 1.66), *p* < 0.001); birth date: (β = -0.72 (95% CI -1.01, -0.46), *p* < 0.001).

In yearlings, in contrast, we uncovered strong negative relationships between survival and local density (β = -0.45 (95% CI -0.78, -0.11), *p* = 0.01, Fig. 2c) and NDVI (β = 0.38 (95% CI 0.07, 0.74), *p* = 0.03, Fig. 2d), but no interactions (*p* > 0.05; ΔDIC < 2, Fig. 4). Consistent with previous studies in this population, males (β = -1.79 (95% CI -2.56, -1.23), *p* < 0.001), those with older mothers (β = -0.50 (95% CI -0.79, -0.23), *p* = 0.004), and those with higher parasite counts (β = -0.79 (95% CI -1.15, -0.45), *p* < 0.001) were less likely to survive. We did not find interactions between parasite counts and calving traits (sex, birth date, birth weight, maternal age) in either calves or yearlings.

For adult female survival, we found a strong negative relationship with local density (β = -0.30 (95% CI -0.52, -0.06), *p* = 0.04), and we confirmed previously-observed costs of raising a calf to winter (β = -1.43 (95% CI -2.04, -0.82), *p* < 0.001), age (β = -1.61 (95% CI -1.84, - 1.32), *p* < 0.001), and parasite count (β = -1.28 (95% CI -1.81, -0.80), *p* < 0.001). There was also a positive interaction between reproduction (raising a calf to winter) and parasite count. That is, there were stronger survival costs of parasitism in non-reproductive than reproductive females (β = 0.92 (95% CI 0.43, 1.45), *p* < 0.001).

Female fecundity was likewise negatively-influenced by raising a calf to winter (β = -4.31 (95% CI -4.92, -3.71), *p* < 0.001), age (β = -0.45 (95% CI -0.77, -0.23), *p* = 0.002), and parasite count (β = -0.26 (95% CI -0.44, -0.03), *p* = 0.01) as previously noted. We also uncovered a strong positive relationship between NDVI and fecundity (β = 0.51 (95% CI 0.20, 0.78), *p* = 0.001), and a strong positive interaction between density and parasite count (β = 0.22 (95% CI 0.03, 0.41), *p* = 0.02). That is, there were stronger fecundity costs of parasitism in females in lower-density than in higher-density areas.

## Discussion

We uncovered pervasive relationships between fine-scale local density, resource availability, parasitism, and fitness in individual wild red deer on the Isle of Rum. All measures of density and resource availability influenced at least one fitness component, and several influenced the strength of parasites’ fitness costs. Density and resource scarcity exacerbated parasites’ survival costs in calves as expected, while adults showed counterintuitive interaction patterns: adults with greater resource demands (i.e., those that reproduced or occupied higher-density areas) exhibited weaker costs of parasites for survival and fecundity respectively. As such, our results imply that parasites and resource scarcity enact synergistic density-dependent population regulation in calves, which fades in yearlings and then inverts itself in adults. Nevertheless, main relationships between fitness and density, resources, and parasites remained in yearlings and adults, indicating that these factors still comprise important selective pressures. We suggest that a selection filter acts on these synergistic costs of parasites and density, leaving behind certain individuals and in certain areas that then do not show the same intensified trends that calves do. As such, considering the spatiotemporal legacy of parasite- and resource-mediated extrinsic mortality is likely to be important when examining population regulation in wild animals.

As expected, high local density intensified parasites’ survival costs in calves, while higher NDVI (i.e., greater resource availability) buffered these negative relationships. Resources are the most likely mediator, supported by the fact that density was no longer significant when NDVI was incorporated, and by their almost perfectly inverted prediction surfaces (Fig. 2a-b). Mechanistically, better nutrition in higher-NDVI areas likely conveyed improved tolerance to heavy strongyle infection, allowing residents to heal or prevent damage, whereas the reverse was true in high-density, low-resource areas. While it is commonly theorized that abundant resources can provide relief from costs of parasitism, most evidence comes from lab studies such as mouse malaria^19^ or gut worms in toads^58^. Limited evidence is available from field studies of wild animals. McNew et al.^59^ used two years of data on mockingbirds in the Galápagos to show that tolerance of nestlings to nest parasites decreased when resources were scarce in years with low rainfall, while Agostini et al.^60^ showed that the severity of parasite fitness costs (i.e., reduction in the ability of the host to forage) only manifested in wild capuchins that did not have access to food provisioning. This is to our knowledge the first time this has been demonstrated in a wild mammal, and the first time that density has been implicated in mediating relationships between resource availability, parasitism, and fitness. Future studies could explore the expression of gastrointestinal immune factors that mediate tolerance to investigate the role of immunological tolerance in this system and its differential expression across space.

Regardless of the precise cause, there are likely to be complex epidemiological implications of density undermining tolerance as well as increasing exposure. Most fundamentally, parasites will exhibit disproportionately strong fitness costs at high densities, amplifying density-dependent mortality. This phenomenon could strengthen parasite-mediated population regulation beyond the expectations of theoretical models that do not consider density-dependent survival costs, some of which have important consequences for conservation planning^61-64^. Negative density-tolerance relationships could also affect evolutionary outcomes. For example, tolerance itself is thought to become more advantageous (relative to resistance) with greater exposure rates, which produces further positive feedback that drives greater parasite prevalence^33,35^. If tolerance and exposure rates are negatively correlated (e.g., via density), this could undermine the benefits of inducing tolerance, undermining this positive feedback cycle. Ultimately this would select against the evolution and expression of tolerance; however, in the absence of an advanced eco-evolutionary model (such as^65-67^), it is unclear how these dynamics might manifest, and how severe the density-dependent undermining of tolerance would need to be. More broadly, several studies have modeled tolerance evolution in a spatially-explicit context, finding that its expression should depend on the presence of spatial structuring^68^. By linking tolerance with spatially-varying factors (density and NDVI), we provide evidence that this is the case. Implications are varied, but they include the possibility that tolerance might be kin-structured and dependent on patterns of conspecifics^68,69^.

As expected, we also found strong relationships between global density and fitness in calves^70^, alongside newly-discovered relationships between local density and NDVI and survival in yearlings (with no relationship with global density). These observations, and specifically the ability of these local density and NDVI metrics to identify these novel relationships, strongly supports the value of using within-population, within-year spatial metrics in addition to annual measures of population size when investigating relationships between resource availability, density, and fitness. There were surprisingly no main relationships between survival and these metrics in calves, which indicates that this fitness component is indeed more dependent on global competition and resource availability at the population level rather than the local level.

In adult females, relationships between parasites and fitness were also strongly modified by ecological context, but counterintuitively. Specifically, survival costs of parasitism were stronger in non-reproductive females than in those raising a calf to winter, which is contrary to our expectation. One possible explanation is that this pattern reflects variation in individual quality: females capable of reproducing while infected may represent higher-quality individuals better able to tolerate parasite burdens, while individuals in poor condition are both less likely to reproduce and more susceptible to parasite-associated mortality. This uncertainty is part of an inherent difficulty extricating causality in observational systems like ours, but our data are nevertheless able to inform the underlying processes. A similarly context-dependent pattern emerged for fecundity, where costs of parasitism were surprisingly strongest in females occupying *lower*-density areas. This counterintuitive pattern may again reflect variation in individual quality or spatial structuring, where higher-density areas disproportionately comprise higher-quality individuals due to the competitive purging of weaker individuals. This trend could arise from the patterns of shooting in the population, where individuals inhabiting the lower-density edges regularly stray outside the study area and risk being shot. As an anthropogenic filter, this source of extrinsic mortality is less tied to individuals’ quality, resources, or parasite burden, and was not examined here: shot animals were removed from the data and not analyzed. Given that parasites are often observed to affect vigilance, movement, and other components of predation risk^71-73^, it is possible parasites also have an effect on the probability of being shot, and that this is influencing our results. One analysis showed that *Toxoplasma gondii*-infected red deer were culled more quickly than uninfected counterparts under a stringent set of conditions^74^, but another study shows no difference in hunting rates of white-tailed deer based on whether they were infected with chronic wasting disease^75^. Future studies could test these interrelationships in this system and their role in determining patterns of mortality and space use.

Together, the juvenile and adult results demonstrate the importance of considering life stage and spatiotemporal demographic structuring when evaluating the fitness costs of parasitism. In calves, parasites interacted with increased local density and resource scarcity, whereas these relationships largely disappeared or reversed in older individuals. The best explanation for this disparity is that selective processes progressively filter individuals as they age: individuals most susceptible to parasite-associated mortality are removed early in life, leaving a subset of more robust individuals in older age classes. Consequently, parasite-fitness relationships observed in adults may not reflect the full spectrum of ecological interactions present earlier in life. This pattern suggests that early life stages are likely particularly valuable when investigating how ecological conditions shape parasite costs. Young animals often show weaker immunity and increased parasite infection, alongside stronger density-dependence, which has produced patterns like the one we report here. Indeed, a study in Soay sheep (*Ovis aries*) showed a strong positive relationship between local density and gastrointestinal parasite counts in juveniles that faded in adults, which is thought to have occurred through a similar filtering^10^. These sorts of age-structured density-dependent parasite costs may be common in wild mammals, or at least in ungulates. Particularly in capital breeding species such as red deer, individuals that reach adulthood may already represent a filtered subset of the population, obscuring density- and resource-mediated parasite effects that are more evident in juveniles.

These findings highlight that parasite-mediated fitness costs depend strongly on ecological context, supporting prior calls for the inclusion of parasites into ecological and evolutionary frameworks^76-80^. This paper adds to the literature on density-dependent trends in parasite infection, expanding them from merely identifying density-infection trends to demonstrating that they cause fitness costs to vary in severity. Although parasites are often assumed to regulate populations, our findings show that these relationships can vary substantially at fine-scale spatiotemporal scales and across life stages. Understanding how parasites influence population processes therefore requires considering not only infections but also the place and time in which they occur. That is, parasites do not just determine who lives and dies, but who lives and dies *where*.

## Acknowledgements

We thank NatureScot for permission to work on the Isle of Rum and the many volunteers and researchers who have helped at the field site during this study. We thank Dave McBean, Moredun Institute, for some of the FECs. JMP, GFA, AZH, and the Rum deer project acknowledge funding from the Leverhulme Trust (RPG 2022-220). GFA acknowledges funding from WAI (C-2023-00057). AZH benefitted from the musical inspiration of Unto Others.

