## Supplemental materials for "Fitness costs of parasitism depend on fine-scale density and resource availability in a wild ungulate"

**Additional details of NDVI calculations**

We used the 'LandsatTS' package^1^ to acquire Google Earth Engine-hosted Level-2 Collection-2 Tier-1 Landsat 5 (Thematic Mapper [TM]), Landsat 7 (Enhanced Thematic Mapper Plus [ETM+]), and Landsat 8 (Operational Land Imager and Thermal Infra-Red Scanner [OLI-TIRS]) satellite imagery. Throughout the study period, approximately one image of the study area was acquired per week, each with a pixel-level resolution of 30 meters. Each image was pre-processed to normalize surface reflectance and to categorize each pixel using the automated function of mask (cfmask) algorithm^2^. Prior to analysis, we excluded all pixels categorized as no data, cloud, cloud shadow, snow, or water, ensuring that only 'clear' ground surface pixels were used to calculate NDVI. We limited the dataset to pixels containing NDVI values of 0.15 and above, as values below this threshold typically indicate non-biomass areas such as concrete or buildings. Additionally, due to systematic differences in surface reflectance and spectral indices among Landsat sensors^3^, we cross-calibrated the data among sensors to ensure that NDVI values were comparable regardless of the sensor. Cross-calibration followed the random forest model workflow of Berner et al.^1^ (available from https://github.com/logan-berner/LandsatTS). In brief, the approach involved identifying the characteristic reflectance at sample sites during the growing season, defined as the beginning of March to the end of September, using Landsat 7 and Landsat 5/8 data from the same years. This was used this to train a random forest model to predict Landsat 5/8 reflectance from the Landsat 7 reflectance values. To account for a dearth of valid NDVI pixel data with which to train the model, we employed the high-latitude training dataset provided in the LandsatTS package. From these data, we again used a process from LandsatTS package by Berner et al.^1^ to quantify the growing season characteristics. This involved iteratively fitting cubic splines to pixel measurements pooled over a seven-year moving window within the growing season. Observations were exponentially-weighted by distance in years from the focal year, so that observations from the focal year were most important in calculating its spline. Outliers were excluded and the splines refitted until all observations were within a 30% bound of the fitted spline. If there were fewer than ten observations in the focal window, the spline was not fit. We then estimated the maximum NDVI and the associated day of year for each pixel from its fitted spline. We calculated mean annual NDVI values (hereafter NDVI) for the deer by averaging the NDVI values from each sighting of each deer throughout a given year, giving us an estimate of the maximum amount of vegetation a given deer had access to during that year, giving us an estimate of the maximum amount of vegetation a given deer had access to during a given year.
